# Predictive coding with neural transmission delays: a real-time temporal alignment hypothesis

**DOI:** 10.1101/453183

**Authors:** Hinze Hogendoorn, Anthony N Burkitt

## Abstract

Hierarchical predictive coding is an influential model of cortical organization, in which sequential hierarchical layers are connected by feedback connections carrying predictions, as well as feedforward connections carrying prediction errors. To date, however, predictive coding models have neglected to take into account that neural transmission itself takes time. For a time-varying stimulus, such as a moving object, this means that feedback predictions become misaligned with new sensory input. We present an extended model implementing both feed-forward and feedback extrapolation mechanisms that realigns feedback predictions to minimize prediction error. This realignment has the consequence that neural representations across all hierarchical stages become aligned in real-time. Using visual motion as an example, we show that the model is neurally plausible, that it is consistent with evidence of extrapolation mechanisms throughout the visual hierarchy, that it predicts several known motion-position illusions, and that it provides a solution to the temporal binding problem.

## Introduction

Predictive coding is a model of neural organisation that originates from a long history of proposals that the brain infers, or predicts, the state of the world on the basis of sensory input ^1–3^. It has been particularly influential in the domain of visual perception ^4–8^, but has also been extensively applied in audition (see ^9,10^ for reviews), somatosensation (e.g. ^11^), motor control ^12,13^, and decision science ^14,15^, where it accounts for a range of subtle response properties and accords with physiology and neuroanatomy. Although it has been criticized for being insufficiently articulated^16^, it has been further developed into a general theory of cortical organization^17,18^ and even been advocated as a fundamental principle of self-organising biological systems in general (the Free Energy Principle, ^19–22^).

An essential principle of predictive coding is a functional organization in which higher organizational units “predict” the activation of lower units. Those lower units then compare their afferent input to this feedback prediction, and feed forward the difference: a prediction error^4^ (where the term “prediction” is used in the strictly hierarchical sense, rather than the everyday temporal sense of predicting the future). This interaction of feedback predictions and feedforward prediction errors characterizes each subsequent level of the processing hierarchy (Figure 1A).

**Figure 1:**
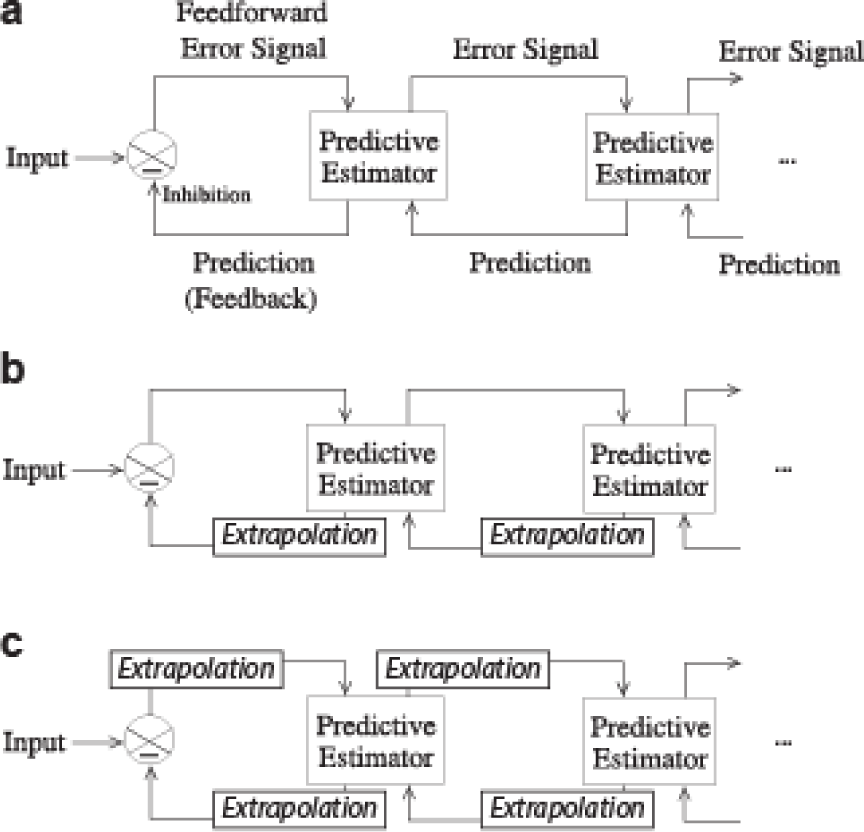
*The classical hierarchical predictive coding model and two possible extensions. **[a]** The **Classical Predictive Model**. This model consists of hierarchically organized loops of feedforward and feedback connections. Feedback signals carry predictions, and feedforward signals carry prediction errors.^4^ **[b]** The **Predictive Model with Extrapolated Feedback**. In order to handle time-varying stimuli such as motion, the classical model can be expanded to include an extrapolation mechanism on the feedback step. This would be one way to minimize prediction error for time-varying stimuli. **[c]** The **Predictive Model with Real-Time Alignment**. In this model, extrapolation mechanisms work on both feedforward and feedback steps. Like model (b), this would minimize prediction error, but has the additional consequence that it aligns the content of neural representations across the hierarchy. Diagram labels in (b) and (c) are as in (a) but omitted for clarity*.

In this review, we argue that neural transmission delays cause the classical model of hierarchical predictive coding^4^ to break down when input to the hierarchy changes on a timescale comparable with that of the neural processing. Using visual motion as an example, we present two models that extend the classical model with extrapolation mechanisms to minimize prediction error for time-varying sensory input. One of these models, in which extrapolation mechanisms operate on both feedforward and feedback mechanisms, has as a consequence that all stages in the hierarchy become aligned in real-time. We argue that this model not only minimizes prediction error, but also parsimoniously explains a range of spatiotemporal phenomena, including motion-induced position shifts such as the flash-lag effect and related illusions, and provides a natural solution to the question how asynchronous processing of visual features nevertheless leads to a synchronous conscious experience (the temporal binding problem).

## Minimizing Prediction Error

The core principle behind the pattern of functional connectivity in hierarchical predictive coding is the minimization of total prediction error. This is considered to be the driving principle on multiple biological time-scales ^22^:

At long time-scales, the minimization of prediction error drives evolution and phylogenesis. Neural signaling is metabolically expensive, and there is therefore evolutionary pressure for an organism to evolve a system of neural representation that allows for complex patterns of information to be represented with minimal neural firing. A sparse, higher-order representation that inhibits the costly, redundant activity of lower levels would provide a metabolic, and therefore evolutionary, advantage.

At the level of individual organisms, at time-scales relevant to decision-making and behaviour, the minimization of prediction error drives learning ^22,23^. Predictions are made on the basis of an internal model of the world, with the brain essentially using sensory input to predict the underlying cause(s) of that input. The better this internal model fits the world, the better the prediction that can be made, and the lower the prediction error. Minimizing prediction error therefore drives a neural circuit to improve its representation of the world, in other words: learning.

Finally, at sub-second time-scales relevant to sensory processing, the minimization of prediction error drives the generation of stable perceptual representations. A given pattern of sensory input feeds in to feedforward and feedback mechanisms that iteratively project to each other until a dynamic equilibrium is reached between higher-order predictions and (local) deviations from those predictions. Because this equilibrium is the most efficient representation of the incoming sensory input, the principle of prediction error minimization works to maintain this representation as long as the stimulus is present. This results in a perceptual representation that remains stable over time. Interestingly, because there may be local minima in the function determining total prediction error, a given stimulus might have multiple stable interpretations, as is for example the case for ambiguous stimuli such as the famous Necker cube ^24^.

## Hierarchical vs Temporal Prediction

In discussing predictive coding, there is an important distinction to be made regarding the sense in which predictive coding models predict.

Descriptively, predictive coding models are typically considered (either implicitly or explicitly) to reflect some kind of expectation about the *future*. For example, in perception, pre-activation of the neural representation of an expected sensory event ahead of that event’s actual occurrence reflects the nervous system predicting a *future* event ^10,25,26^. In a decision-making context, a prediction might likewise be a belief about the future consequences of a particular choice ^27^. These are predictions in the temporal sense of predicting future patterns of neural activity.

However, mechanistic models of predictive coding such as the one first proposed by Rao and Ballard are not predictive in this same sense^4^. Rather than predicting the future, these models are predictive in the *hierarchical* sense of higher areas predicting the activity of lower areas ^4,8,17,18^. These models do not predict what *is going to happen*: rather, by converging on a configuration of neural activity that optimizes the representation of a stable sensory input, they hierarchically predict *what is happening*. The use of the word “prediction” in this context is therefore somewhat unfortunate, as mechanistic models of predictive coding do not actually present a mechanism that predicts in the temporal way that “prediction” is typically used in ordinary discourse.

## Predicting the Future

To date, the temporal dimension has been nearly absent from computational work on mechanistic models of predictive coding. In general, it is implicitly assumed that sensory input remains unchanged until the system converges on a minimal-prediction-error solution. The available studies of dynamic stimuli in a predictive coding context consider only autocorrelations within a given neuron’s time-series – in other words, the tendency of sensory input to remain the same from moment to moment ^28–30^. Computationally, this prediction is easily implemented as a biphasic temporal impulse response function ^29,31,32^, which is consistent with known properties of neurons in the early visual pathway, including the lateral geniculate nucleus (LGN;^31,32^). However, this is only the most minimal temporal prediction that a neural system might make: the prediction that its input remains unchanged over time.

Importantly, what is missing from mechanistic models of predictive coding is the fact that *neural communication itself takes time*. Perhaps because the delays involved in *in silico* simulations are negligible, or perhaps because the model was first articulated for static images and stable neural representations, models of neural circuitry have entirely neglected to take into account that neural transmission incurs significant delays. These delays mean that feedforward and feedback signals are misaligned in time. For an event at a given moment, the sensory representation at the first stage of the hierarchy needs to be fed forward to the next hierarchical level, where a prediction is formulated, which is then fed back to the original level. In the case of a dynamic stimulus, however, by the time that the prediction arrives back at the first hierarchical stage, the stimulus will have changed, and the first stage will be representing the new state of that stimulus. As a result, feedback predictions will be compared against more recent sensory information than the information on which they were originally based, and which they were sent to suppress. If this temporal misalignment between feedforward and feedback signals would not somehow be compensated, under the classical hierarchical predictive coding model any time-varying stimulus would generate very large prediction errors at each level of representation, which is typically not seen in electrophysiological recordings of responses to stimuli with constant motion (as opposed to unexpected changes in the trajectory, e.g. ^33–35^).

Due to neural transmission delays, prediction error is minimized not when a feedback signal represents the sensory information that originally generated it, but when it represents the sensory information that *is going to be* available at the lower level by the time the feedback signal arrives. In other words, prediction error is minimized when the feedback signal *anticipates* the future state of the lower hierarchical stage. Estimating that future state requires only rate-of-change information about the relevant feature, and it follows that if such information is available at a given stage, it will be recruited to minimize prediction error. When allowing for transmission delays, hierarchical predictions therefore need to become temporal predictions: they need to predict the future.

## Two Extended Models

A clear example of common, time-varying sensory input is visual motion. Here, we use visual motion to illustrate the limitations of the classical predictive coding model when input is time-varying, and present two extensions to the classical model that would solve these limitations.

A. **The Classical Predictive Model:** In the classical hierarchical predictive coding model^4^ (Figure 1A), neural transmission delays mean that feedback predictions arrive at an area a significant interval of time after that area fed forward the signal that led to those predictions. Because the input to this level has changed during the elapsed time, this results in prediction error (Figure 2A).
B. **The Predictive Model with Extrapolated Feedback:** In this extension to the classical model, each complete feed-forward/feed-back loop uses any available rate-of-change information to minimize prediction error. In the case of visual motion, this means the circuit will use concurrent velocity information to anticipate the representation at the lower level *by the time the feed-back signal arrives at that level*. In this model, an extrapolation mechanism is implemented at the feedback step of each loop (Figure 1B) to minimize predictive error whilst leaving the feedforward sweep of information unchanged (Figure 2B).
C. **The Predictive Model with Real-Time Alignment:** In the classical model, prediction error results from the cumulative delay of both feedforward and feedback transmission. Prediction error is minimized when this cumulative delay is compensated at *any* point(s) in the feedforward-feedback loop. Evidence from both perception^36,37^ and action^38,39^ strongly suggests that at least part of the total delay incurred is compensated by extrapolation at the feedforward step. Accordingly, in this model, we propose that extrapolation mechanisms work on both feedforward and feedback signals: *any* signal that is sent from one stage to another, whether forward or backward, is extrapolated into the future that precisely compensates for the delay incurred by that signal while it is in transit (Figure 1C). In addition to minimizing prediction error, this model has the remarkable consequence of synchronizing representations throughout the hierarchy: under this model, all areas represent a moving object in the same position at the same moment (Figure 2C), independent of where in the hierarchy each area lies.

**Figure 2:**
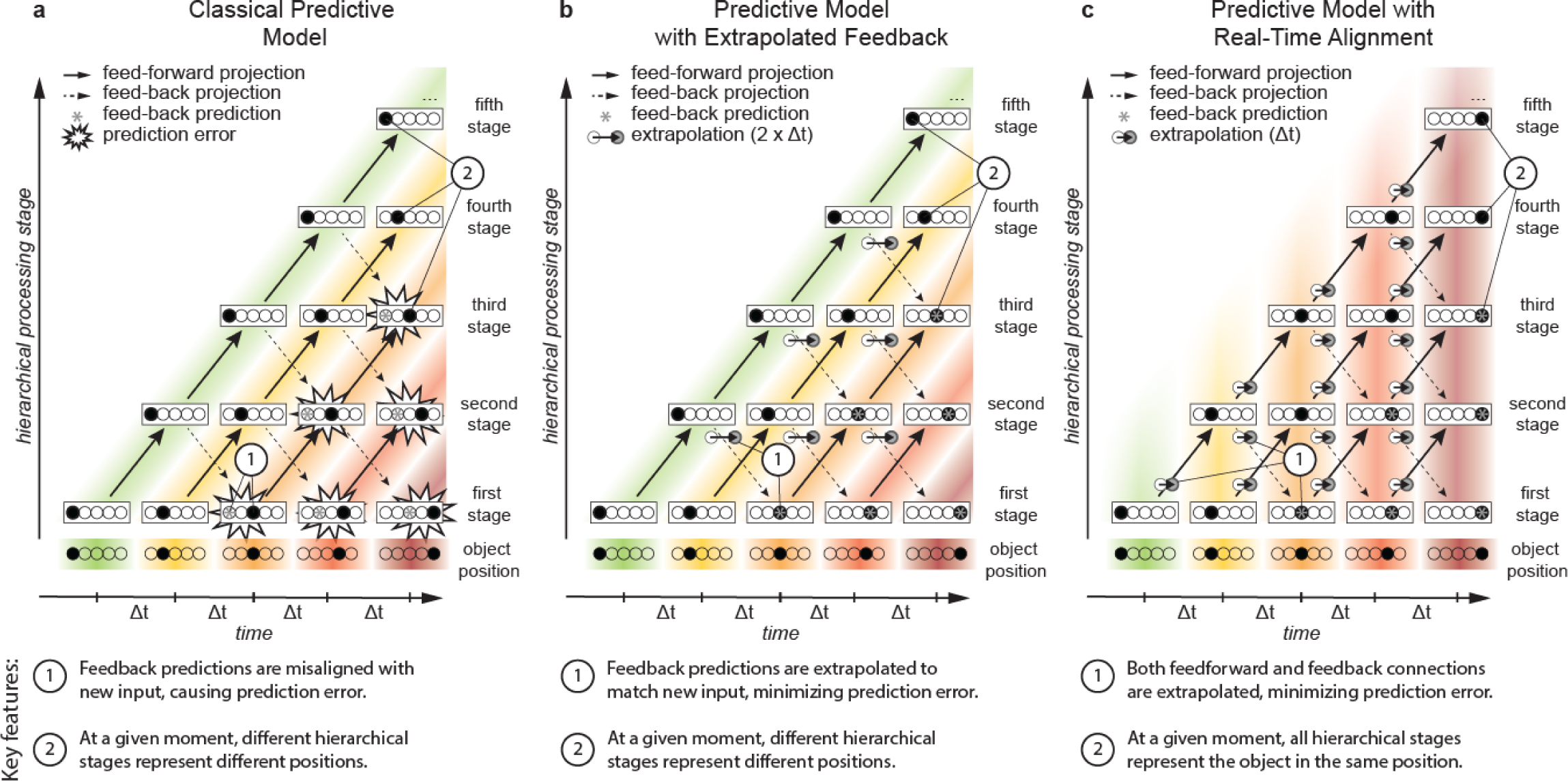
*Three simplified models of the representation of the position of a moving object throughout the visual processing hierarchy under the predictive coding framework. Each rectangle denotes the neural representation of the object’s position at a given hierarchical level and at a given time, with the filled circle indicating the object in one of five possible positions. In this simplified representation, all connections are modeled as incurring an equal transmission delay Δt. Coloured bands link corresponding representations and numbered circles highlight core features of each model **[a].** The **Classical Predictive Model** of predictive coding comprises feedforward connections from one hierarchical level to the next level (solid lines), and feedback connections to the previous level (dashed lines). No allowance is made for neural transmission delays, such that feedback connections carry a position-representation (asterisks) that is outdated by the time that signal arrives. The resulting mismatch with more recent sensory input generates large errors, which subsequently propagate through the hierarchy (emphasized with starbursts). **[b].** In the **Predictive Model with Extrapolated Feedback**, an extrapolation mechanism operates on predictive feed-back projections, anticipating the future position of the object. This mechanism compensates for the total time-cost incurred during both the feed-forward and the feed-back portion of the loop. This would minimize total prediction error in this simplified model. However, the mechanism would rapidly become more complex when one considers that individual areas tend to send and receive signals to and from multiple levels of the hierarchy. **[c].** In the **Predictive Model with Real-Time Alignment**, extrapolation mechanisms compensate for neural delays at both feed-forward and feed-back steps. This parsimoniously minimizes total prediction error, even for more complex connectivity patterns. Additionally, the model differs from the first two models in that at any given time, all hierarchical stages represent the same position. Conversely, in the first two models, at any given time neural transmission delays mean that all hierarchical levels represent a different position. This crucial difference is evident as vertical, rather than diagonal coloured bands linking matching representations across the hierarchy. The consequence of this hypothesis is therefore that the entire visual hierarchy becomes synchronized. This provides an automatic and elegant solution to the computational challenge of establishing which neural signals belong together in time: the temporal binding problem. It is also consistent with demonstrated extrapolation mechanisms in feed-forward pathways and provides a parsimonious explanation for a range of motion-induced position shifts*.

## Evaluating the evidence

We have argued above that the classical model of predictive coding ^4,17,18,29^ will consistently produce prediction errors when stimuli are time-varying and neural transmission delays are taken into consideration (Figure 2A). We have proposed two possible extensions to the classical model, each of which would minimize prediction error. Here, we evaluate the evidence for and against each of these two models.

### Neural Plausibility

Models B and C are both neurally plausible. Both extensions to the classical model incorporate neural extrapolation mechanisms at each stage of the visual hierarchy. This requires firstly that information about rate-of-change be represented and available at each stage. It is well-established that velocity is explicitly represented throughout the early visual hierarchy, including the retina ^40^, LGN ^41–44^, and primary visual cortex ^45^. As required by both models, velocity information is therefore available throughout the hierarchy. Indeed, it follows from the error minimization principle that if velocity is available at each stage, it can and will be used at each stage to optimize the accuracy of feedback predictions involving that stage.

Secondly, the extrapolation processes posited in both models should cause activity ahead of a moving object’s position, preactivating the area of space into which the object is expected to move. This activation ahead of a moving object is consistent with reported preactivation of predicted stimulus position in cat V1^46^, as well as in human EEG^26^. Importantly, if the object unexpectedly vanishes, such extrapolation would pre-activate areas of space in which the object ultimately never appeared. This is consistent with psychophysical experiments where an object is perceived to move into areas of the retina where no actual stimulus energy can be detected^47,48^, as well as with recent fMRI work showing activation in retinotopic areas of the visual field beyond the end of the motion trajectory^49^.

### Hierarchical Complexity

Model C is more robust to hierarchical complexity than Model B. Figure 2 shows extremely simplified hierarchies: it omits connections that span multiple hierarchical levels, it represents all time-delays as equal and constant, and it shows only a single projection from each area. In reality, of course, any given area is connected to numerous other areas^50^, and transmission delays will inevitably differ depending on where feedforward signals are sent to and where feedback signals originate from. Importantly, a given area might receive input about the same moment through different pathways with different lags. A purely feedback extrapolation process (Model B) would not be able to compensate for these different lags, because it would not be able to differentiate between the feedforward signals that led to the prediction. Conversely, a model with extrapolation mechanisms at both feedforward and feedback connections (Model C) *would* be able to compensate for any degree of hierarchical complexity, as each connection would essentially compensate for its own delay.

### Motion-induced Position Shifts

Model C predicts motion-induced position shifts: visual illusions in which motion signals affect the perceived positions of objects. In Model C, the representation in higher hierarchical levels does *not* represent the properties of the stimulus as it was when it was initially detected, but rather the *expected* properties of that stimulus a certain extent into the future. In the case of visual motion, this means that the feedforward signal represents the extrapolated, rather than detected position of the moving object. One consequence is that at higher hierarchical levels, a moving object is never represented in the position at which it first appears (Figure 2C: top diagonal). Instead, it is first represented a certain extent along its (expected) trajectory. Model C therefore neatly predicts a visual phenomenon known as the Fröhlich effect, in which the initial position of an object that suddenly appears in motion is perceived as being shifted in the direction of its motion^51,52^.

Model C is also consistent with the much-studied flash-lag phenomenon^37^, in which a briefly flashed stimulus that is presented in alignment with a moving stimulus appears to lag behind that stimulus. Although alternative explanations have been proposed^53^, a prominent interpretation of this effect is that it reflects motion extrapolation, specifically implemented to compensate for neural delays ^36,37,54^. Model C is not only compatible with that interpretation, but it provides a principled argument (prediction error minimization) for why such mechanisms might develop.

### Neural Computations

Both Model B and C posit interactions between motion and position signals at multiple levels in the hierarchy. This is compatible with a number of theoretical and computational models of motion and position perception. For example, Eagleman and Sejnowski have argued that for a whole class of motion-induced position shifts, a local motion signal biases perceived position^55^ – precisely the local interactions between velocity and position signals that we propose. Furthermore, these instantaneous velocity signals have been shown to affect not only the perceptual localization of concurrently presented targets, but also the planning of saccadic eye movements aimed at those targets^56,57^. Recent computational work has argued that the flash-lag effect reflects a Bayesian integration of motion and position signals, and postulated that such mechanisms might compensate for neural delays^58^. In a similar vein, Kwon and colleagues recently advanced an elegant computational model uniting motion and position perception, and although not formulated in terms of extrapolation, an interaction between motion and position signals was a central premise of their model^59^. We argue that, under the hierarchical predictive coding framework, neural processing delays necessarily lead to the evolution of motion-position interactions.

Furthermore, both Models B and C posit such interactions at multiple stages, including very early stages in the hierarchy. This is consistent with recent work on the flash-grab effect: a motion-induced position shift in which a target briefly flashed on a moving background that reverses direction is perceived shifted away from its true position^60^. Using EEG, the interaction between motion and position signals that generates the illusion has been shown to occur already within the first 80 ms following stimulus presentation, indicating a very early locus of interaction. A follow-up study using dichoptic presentation revealed that even within this narrow time-frame, extrapolation took place at at least two dissociable processing stages^61^.

### Temporal Alignment

A defining feature of Model C is that, due to extrapolation at each feedforward step, all hierarchical areas become aligned in time. Although neural transmission delays mean that it takes successively longer for a new stimulus to be represented at successive layers of the hierarchy^62^, the fact that the neural signal is extrapolated into the future at each stage means that the *representational content* of each consecutive stage now runs aligned with the first hierarchical stage, and potentially with the world, in real time. In the case of motion, we perceive moving objects where they are, rather than where they were, because the visual system extrapolates the position of those objects to compensate for the delays incurred during processing. Of course, the proposal that the brain compensates neural delays by extrapolation is not new (see ^36^ for a review). Rather, what is new here is the recognition that this mechanism has not developed for the *purpose* of compensating for delays at the behavioural level, but that it follows necessarily from the fundamental principles of predictive coding.

The temporal alignment characterizing Model C also provides a natural solution to the problem of temporal binding. Different visual features (such as contours, colour, motion, etc.) are processed in different, specialized brain areas ^50,63^. Due to anatomical and physiological differences, these areas process their information with different latencies, which leads to asynchrony in visual processing. For example, the extraction of colour has been argued to lead the extraction of motion ^64–66^. The question therefore arises how these asynchronously processed features are nevertheless experienced as a coherent stream of visual awareness, with components of that experience processed in different places at different times^67^. Model C intrinsically solves this problem. Because the representation in each area is extrapolated to compensate for the delay incurred in connecting to that area, representations across areas (and therefore features) become aligned in time.

### Local Time Perception

Model C eliminates the need for a central timing mechanism. Because temporal alignment in this model is an automatic consequence of prediction error minimization at the level of local circuits, no central timing mechanism (e.g., internal clock^68^) is required to carry out this alignment. Indeed, under this model, if a temporal property of the prediction loop changed for a particular part of the visual field, then this would be expected to result in *localized* changes in temporal alignment. Although this is at odds with our intuitive experience of the unified passage of time, local adaptation to flicker has been found to distort time perception in a specific location in the visual field^69,70^. Indeed, these spatially localized distortions have been argued to result from adaptive changes to the temporal impulse response function in the lateral geniculate nucleus LGN^71,72^, which would disrupt the calibration of any neuron extrapolating a given amount of time into the future. The counterintuitive empirical finding that the perceived timing of events in specific positions in the visual field can be distorted by local adaptation is therefore consistent with the local compensation mechanisms that form part of Model C.

### Violated Predictions

Models B and C only produce veridical percepts when objects move on predictable trajectories. When events do not unfold predictably, such as when objects unexpectedly appear, disappear, or change their trajectory, this introduces new feedforward information which is, of course, at odds with feedback predictions at each stage. This gives rise to prediction errors at each stage of the hierarchy, with prediction errors getting larger further up the hierarchy (as it takes longer for sensory information signaling a violation of the status quo to arrive at higher stages). During the intervening time, those areas will continue to (erroneously) extrapolate into a future that never comes to pass.

The breakdown of error minimization (and therefore accurate perception) in situations where events unfold unpredictably fits with empirical studies showing visual illusions in these situations. As noted above, the flash-lag effect is one such visual illusion. Under Model B or C, this effect occurs because the position of the moving object can be extrapolated because it is predictable, whereas the flashed object, which is unpredictable, cannot^37^. This mismatch between the predictable motion and the unpredictable flash has a parallel in the flash-grab effect: a recent study parametrically varied the predictability of the flash, and showed that the strength of the resulting illusion decreased as the flash became more predictable^73^. Likewise, in studies of temporal binding, the asynchrony in the processing of colour and motion is evident only when rapidly moving stimuli abruptly change direction^74^. These illusions reveal that accurate perception breaks down when prediction is not possible, consistent with both models B and C, but not with the classical predictive model A.

## Conclusions and Future Directions

Altogether, multiple lines of evidence converge in support of Model C: an extension of the hierarchical predictive coding framework in which extrapolation mechanisms work on both feedforward and feedback connections to minimize prediction error. In this model, minimal prediction error is achieved by local extrapolation mechanisms compensating the specific delays incurred by individual connections at each level of the processing hierarchy. As a result, neural representations across the hierarchy become aligned in real-time. This model provides an extension to classical predictive coding models that is necessary to account for neural transmission delays. In addition, the model predicts and explains a wide range of spatiotemporal phenomena, including motion-induced position shifts, the temporal binding problem, and localized distortions of perceived time.

### Neural Implementation

We have taken error minimization as the organizing principle of predictive coding, but now extended to incorporate the local velocity of the stimuli. This is in keeping with the approach of previous authors, who have successfully modeled hierarchical error minimization ^4,17,18,21,29,75^ and motion-position interactions^59^ as a Kalman filter^76^. The extension proposed here can, in principle, be implemented straight forwardly by incorporating the local velocity as an additional state variable in a manner analogous to that proposed for motion-position interactions^59^. Such an approach, in which the prediction error minimization incorporates the expected changes that occur due to both the motion of the stimulus and the propagation of the neural signals, is possible when the local velocity is one of the state variables that each level of the hierarchy has access to. However, the details of how cortical circuits implement such processing or what synaptic plasticity mechanisms underlie the formation of these circuits still remain to be elaborated, and this is a key area for future research.

### Prediction in the Retina

The proposed model follows from the principle of error minimization within predictive feedback loops. These feedback loops are ubiquitous throughout the visual pathway, including feedback from V1 to LGN. Although there are no feedback connections back to the retina, extrapolation mechanisms have nevertheless been reported in the retina^77^, and these mechanisms have even been found to produce a specific response to reversals of motion direction, much akin to a prediction error^33,78,79^. In the absence of feedback connections to the retina, our model does not directly predict these mechanisms. On the long time-scale, compensation for neural delays in the retina provides a behavioural, and therefore evolutionary, advantage, but more research will be necessary to address whether any short-term learning mechanisms play a role in the development of these circuits.

### The Ubiquity of Velocity Signals

We have emphasized hierarchical mechanisms at early stages of visual processing, consistent with extrapolation in monocular channels^61^, early EEG correlates of extrapolation^80^, and evidence that extrapolation mechanisms are shared for both perceptual localization and saccadic targeting^57^. However, an influential body of literature has proposed that the human visual system is organized into two (partly) dissociable pathways: a ventral “what” pathway for object recognition and identification, and a dorsal “where” pathway for localization and motion perception ^81–84^. Two decades later, the distinction is more nuanced^85,86^, but the question remains whether velocity signals are truly ubiquitous throughout the visual hierarchy. Consequently, whether real-time temporal alignment is restricted to the “where” pathway or is a general feature of cortical processing remains to be elucidated.

### The Functional Role of Prediction Error

Throughout this Perspective, we have considered prediction error as something that an effective predictive coding model should minimize. We have not addressed the functional role of the prediction error signal itself. As noted by several authors, this signal might serve an alerting or surprise function (see ^23^ for a review). In the model proposed here, the signal might have the additional function of correcting a faulty extrapolation ^36,48,54^. In this role, prediction error signals would work to eliminate lingering neural traces of predictions that were unsubstantiated by sensory input: expected events that did not end up happening. This corrective function has been modelled as a “catch-up” effect for trajectory reversals in the retina ^33,78^, and as an increase in position uncertainty (and therefore an increase in relative reliance on sensory information) when objects change trajectory in a recently proposed Bayesian model of visual motion and position perception^59^. Further identifying the functional role of these signals is an exciting avenue for future research.

## Acknowledgments

HH was supported by the Australian Government through the Australian Research Council’s Discovery Projects funding scheme (project DP180102268).

## References

1. Gregory, R. L. Perceptions as hypotheses. Philos. Trans. R. Soc. Lond. B. Biol. Sci. 290, 181–97 (1980).

2. Dayan, P., Hinton, G. E., Neal, R. M. & Zemel, R. S. The Helmholtz machine. Neural Comput. 7, 889–904 (1995).

3. von Helmholtz, H. Handbuch der physiologischen Optik. (Voss, 1867).

4. Rao, R. P. N. & Ballard, D. H. Predictive coding in the visual cortex: A functional interpretation of some extra-classical receptive-field effects. Nat. Neurosci. 2, 79–87 (1999).

5. Mumford, D. On the computational architecture of the neocortex. Biol. Cybern. 66, 241–251 (1992).

6. Srinivasan, M. V, Laughlin, S. B. & Dubs, A. Predictive coding: a fresh view of inhibition in the retina. Proc. R. Soc. London. Ser. B, Biol. Sci. 216, 427–59 (1982).

7. Spratling, M. W. Reconciling Predictive Coding and Biased Competition Models of Cortical Function. Front. Comput. Neurosci. 2, 4 (2008).

8. Spratling, M. W. Predictive coding accounts for V1 response properties recorded using reverse correlation. Biol. Cybern. 106, 37–49 (2012).

9. Bendixen, A., SanMiguel, I. & Schröger, E. Early electrophysiological indicators for predictive processing in audition: A review. Int. J. Psychophysiol. 83, 120–131 (2012).

10. Garrido, M. I., Kilner, J. M., Stephan, K. E. & Friston, K. J. The mismatch negativity: a review of underlying mechanisms. Clin. Neurophysiol. 120, 453–63 (2009).

11. van Ede, F., de Lange, F., Jensen, O. & Maris, E. Orienting attention to an upcoming tactile event involves a spatially and temporally specific modulation of sensorimotor alpha‐ and beta-band oscillations. J. Neurosci. 31, 2016–24 (2011).

12. Blakemore, S.-J., Goodbody, S. J. & Wolpert, D. M. Predicting the consequences of our own actions: the role of sensorimotor context estimation. J. Neurosci. 18, 7511–7518 (1998).

13. Adams, R. A., Shipp, S. & Friston, K. J. Predictions not commands: Active inference in the motor system. Brain Structure and Function 218, 611–643 (2013).

14. Schultz, W. Predictive Reward Signal of Dopamine Neurons. J. Neurophysiol. 80, 1–27 (1998).

15. Summerfield, C. & de Lange, F. P. Expectation in perceptual decision making: neural and computational mechanisms. Nat. Rev. Neurosci. 15, 745–756 (2014).

16. Kogo, N. & Trengove, C. Is predictive coding theory articulated enough to be testable? Front. Comput. Neurosci. 9, 111 (2015).

17. Bastos, A. M. et al. Canonical Microcircuits for Predictive Coding. Neuron 76, 695–711 (2012).

18. Spratling, M. W. A review of predictive coding algorithms. Brain Cogn. 112, 92–97 (2017).

19. Friston, K. & Kiebel, S. Predictive coding under the free-energy principle. Philos. Trans. R. Soc. B Biol. Sci. 364, 1211–1221 (2009).

20. Friston, K. A theory of cortical responses. Philos. Trans. R. Soc. B Biol. Sci. 360, 815–836 (2005).

21. Friston, K. The free-energy principle: a unified brain theory? Nat. Rev. Neurosci. 11, 127–138 (2010).

22. Friston, K. Does predictive coding have a future? Nat. Neurosci. 21, 1019–1021 (2018).

23. den Ouden, H. E. M., Kok, P. & de Lange, F. P. How Prediction Errors Shape Perception, Attention, and Motivation. Front. Psychol. 3, 1–12 (2012).

24. Necker, L. A. LXI. Observations on some remarkable optical phænomena seen in Switzerland; and on an optical phænomenon which occurs on viewing a figure of a crystal or geometrical solid. London, Edinburgh, Dublin Philos. Mag. J. Sci. 1, 329–337 (1832).

25. Kok, P., Mostert, P. & de Lange, F. P. Prior expectations induce prestimulus sensory templates. Proc. Natl. Acad. Sci. U. S. A. 114, 10473–10478 (2017).

26. Hogendoorn, H. & Burkitt, A. N. Predictive coding of visual object position ahead of moving objects revealed by time-resolved EEG decoding. Neuroimage 171, (2018).

27. Sterzer, P., Voss, M., Schlagenhauf, F. & Heinz, A. Decision-making in schizophrenia: A predictive-coding perspective. Neuroimage (2018). doi:10.1016/j.neuroimage.2018.05.074

28. van Hateren, J. H. & Ruderman, D. L. Independent component analysis of natural image sequences yields spatio-temporal filters similar to simple cells in primary visual cortex. Proc. R. Soc. Lond. B 265, 2315–2320 (1998).

29. Huang, Y. & Rao, R. P. N. Predictive coding. Wiley Interdiscip. Rev. Cogn. Sci. 2, 580–593 (2011).

30. Rao, R. P. N. An optimal estimation approach to visual perception and learning. Vision Res. 39, 1963–1989 (1999).

31. Dan, Y., Atick, J. J. & Reid, R. C. Efficient Coding of Natural Scenes in the Lateral Geniculate Nucleus: Experimental Test of a Computational Theory. J. Neurosci. 16, 3351–3362 (1996).

32. Dong, D. W. & Atick, J. J. Temporal decorrelation: A theory of lagged and nonlagged responses in the lateral geniculate nucleus. Netw. Comput. Neural Syst. 6, 159–178 (1995).

33. Schwartz, G. et al. Synchronized firing among retinal ganglion cells signals motion reversal. Neuron 55, 958–69 (2007).

34. McMillan, G. A. & Gray, J. R. A looming-sensitive pathway responds to changes in the trajectory of object motion. J. Neurophysiol. 108, 1052–1068 (2012).

35. Dick, P. C. & Gray, J. R. Spatiotemporal stimulus properties modulate responses to trajectory changes in a locust looming-sensitive pathway. J. Neurophysiol. 111, 1736–1745 (2014).

36. Nijhawan, R. Visual prediction: Psychophysics and neurophysiology of compensation for time delays. Behav. Brain Sci. 31, 179–239 (2008).

37. Nijhawan, R. Motion extrapolation in catching. Nature 370, 256–257 (1994).

38. Zago, M., McIntyre, J., Senot, P. & Lacquaniti, F. Visuo-motor coordination and internal models for object interception. Exp. Brain Res. 192, 571–604 (2009).

39. Soechting, J. F., Juveli, J. Z. & Rao, H. M. Models for the extrapolation of target motion for manual interception. J. Neurophysiol. 102, 1491–502 (2009).

40. Barlow, H. B. & Levick, W. R. The mechanism of directionally selective units in rabbit’s retina. J. Physiol. 178, 477–504 (1965).

41. Cheong, S. K., Tailby, C., Solomon, S. G. & Martin, P. R. Cortical-Like Receptive Fields in the Lateral Geniculate Nucleus of Marmoset Monkeys. J. Neurosci. 33, 6864–6876 (2013).

42. Marshel, J. H., Kaye, A. P., Nauhaus, I. & Callaway, E. M. Anterior-posterior direction opponency in the superficial mouse lateral geniculate nucleus. Neuron 76, 713–20 (2012).

43. Zaltsman, J. B., Heimel, J. A. & Van Hooser, S. D. Weak orientation and direction selectivity in lateral geniculate nucleus representing central vision in the gray squirrel Sciurus carolinensis. J. Neurophysiol. 113, 2987–97 (2015).

44. Cruz-Martín, A. et al. A dedicated circuit linking direction selective retinal ganglion cells to primary visual cortex. Nature 507, 358 (2014).

45. Hubel, D. H. & Wiesel, T. N. Receptive Fields and Functional Architecture of monkey striate cortex. J. Physiol. 195, 215–243 (1968).

46. Jancke, D., Erlhagen, W., Schöner, G. & Dinse, H. R. Shorter latencies for motion trajectories than for flashes in population responses of cat primary visual cortex. J. Physiol. 556, 971–982 (2004).

47. Maus, G. W. & Nijhawan, R. Motion Extrapolation Into the Blind Spot. Psychol. Sci. 19, 1087–1091 (2008).

48. Shi, Z. & Nijhawan, R. Motion extrapolation in the central fovea. PLoS One 7, 33651 (2012).

49. Schellekens, W., van Wezel, R. J. A., Petridou, N., Ramsey, N. F. & Raemaekers, M. Predictive coding for motion stimuli in human early visual cortex. Brain Struct. Funct. 221, 879–890 (2016).

50. Felleman, D. J. & Van Essen, D. C. Distributed hierarchical processing in the primate cerebral cortex. Cereb. Cortex 1, 1–47 (1991).

51. Kirschfeld, K. & Kammer, T. The Fröhlich effect: a consequence of the interaction of visual focal attention and metacontrast. Vision Res. 39, 3702–9 (1999).

52. Fröhlich, F. W. Über die Messung der Empfindungszeit. Pflugers Arch. Gesamte Physiol. Menschen Tiere 202, 566–572 (1924).

53. Eagleman, D. M. & Sejnowski, T. J. Motion integration and postdiction in visual awareness. Science 287, 2036–8 (2000).

54. Nijhawan, R. Neural delays, visual motion and the flash-lag effect. Trends Cogn. Sci. 6, 387 (2002).

55. Eagleman, D. M. & Sejnowski, T. J. Motion signals bias localization judgments: A unified explanation for the flash-lag, flash-drag, flash-jump, and Frohlich illusions. J. Vis. 7, 3–3 (2007).

56. Quinet, J. & Goffart, L. Does the Brain Extrapolate the Position of a Transient Moving Target? J. Neurosci. 35, 11780–11790 (2015).

57. Van Heusden, E., Rolfs, M., Cavanagh, P. & Hogendoorn, H. Motion extrapolation for eye movements predicts perceived motion-induced position shifts. J. Neurosci.

58. Khoei, M. A., Masson, G. S. & Perrinet, L. U. The Flash-Lag Effect as a Motion-Based Predictive Shift. PLOS Comput. Biol. 13, e1005068 (2017).

59. Kwon, O.-S., Tadin, D. & Knill, D. C. Unifying account of visual motion and position perception. Proc. Natl. Acad. Sci. U. S. A. 112, 8142–7 (2015).

60. Cavanagh, P. & Anstis, S. M. The flash grab effect. Vision Res. 91, 8–20 (2013).

61. Van Heusden, E., Harris, A. M., Garrido, M. I. & Hogendoorn, H. Predictive coding of visual motion in both monocular and binocular human visual processing.

62. Lamme, V. A. F., Supèr, H. & Spekreijse, H. Feedforward, horizontal, and feedback processing in the visual cortex. Curr. Opin. Neurobiol. 8, 529–535 (1998).

63. Livingstone, M. & Hubel, D. Segregation of form, color, movement, and depth: Anatomy, physiology, and perception. Science (80‐.). 240, 740–749 (1988).

64. Zeki, S. & Bartels, A. The asynchrony of consciousness. Proceedings. Biol. Sci. 265, 1583–5 (1998).

65. Arnold, D. H., Clifford, C. W.. & Wenderoth, P. Asynchronous processing in vision: Color leads motion. Curr. Biol. 11, 596–600 (2001).

66. Moutoussis, K. & Zeki, S. Functional segregation and temporal hierarchy of the visual perceptive systems. Proceedings. Biol. Sci. 264, 1407–14 (1997).

67. Zeki, S. & Bartels, A. Toward a Theory of Visual Consciousness. Conscious. Cogn. 8, 225–259 (1999).

68. Gibbon, J. Scalar expectancy theory and Weber’s law in animal timing. Psychol. Rev. (1977). doi:10.1037/0033-295X.84.3.279

69. Hogendoorn, H., Verstraten, F. A. J. & Johnston, A. Spatially localized time shifts of the perceptual stream. Front. Psychol. 1, 1–8 (2010).

70. Johnston, A., Arnold, D. H. & Nishida, S. Spatially Localized Distortions of Event Time. Curr. Biol. 16, 472–479 (2006).

71. Johnston, A. Visual Time Perception. in The new visual neurosciences (eds. Werner, J. S. & Chalupa, L. M.) 749–762 (MIT Press, 2014).

72. Johnston, A. Modulation of time perception by visual adaptation. in Attention and Time 187–200 (Oxford University Press, 2010). doi:10.1093/acprof:oso/9780199563456.003.0014

73. Adamian, N. & Cavanagh, P. Localization of flash grab targets is improved with sustained spatial attention. J. Vis. 16, 1266 (2016).

74. Moutoussis, K. & Zeki, S. A direct demonstration of perceptual asynchrony in vision. Proceedings. Biol. Sci. 264, 393–9 (1997).

75. Rao, R. P. N. & Ballard, D. H. Dynamic Model of Visual Recognition Predicts Neural Response Properties in the Visual Cortex. Neural Comput. 9, 721–763 (1997).

76. Kalman, R. E. A New Approach to Linear Filtering and Prediction Problems. J. Basic Eng. 82, 35 (1960).

77. Berry, M. J., Brivanlou, I. H., Jordan, T. A. & Meister, M. Anticipation of moving stimuli by the retina. Nature 398, 334–338 (1999).

78. Holy, T. E. A public confession: the retina trumpets its failed predictions. Neuron 55, 831–2 (2007).

79. Chen, E. Y., Chou, J., Park, J., Schwartz, G. & Berry, M. J. The Neural Circuit Mechanisms Underlying the Retinal Response to Motion Reversal. J. Neurosci. (2014). doi:10.1523/JNEUROSCI.1460-13.2014

80. Hogendoorn, H., Verstraten, F. A. J. & Cavanagh, P. Strikingly rapid neural basis of motion induced position shifts revealed by high temporal-resolution EEG pattern classification. Vision Res. 113, (2015).

81. Mishkin, M. & Ungerleider, L. G. Contribution of striate inputs to the visuospatial functions of parieto-preoccipital cortex in monkeys. Behav. Brain Res. 6, 57–77 (1982).

82. Goodale, M. A., Milner, A. D., Jakobson, L. S. & Carey, D. P. A neurological dissociation between perceiving objects and grasping them. Nature 349, 154–156 (1991).

83. Aglioti, S., DeSouza, J. F. & Goodale, M. A. Size-contrast illusions deceive the eye but not the hand. Curr. Biol. 5, 679–85 (1995).

84. Goodale, M. A. & Milner, A. D. Separate visual pathways for perception and action. Trends Neurosci. 15, 20–5 (1992).

85. Milner, A. D. & Goodale, M. A. Two visual systems re-viewed. Neuropsychologia 46, 774–785 (2008).

86. Gilaie-Dotan, S. Visual motion serves but is not under the purview of the dorsal pathway. Neuropsychologia 89, 378–392 (2016).

